# A non-transcriptional function of Yap orchestrates the DNA replication program

**DOI:** 10.1101/2021.11.18.468628

**Authors:** Rodrigo Meléndez García, Olivier Haccard, Albert Chesneau, Hemalatha Narassimprakash, Jérôme E Roger, Muriel Perron, Kathrin Marheineke, Odile Bronchain

## Abstract

In multicellular eukaryotic organisms, the initiation of DNA replication occurs asynchronously throughout S-phase according to a regulated replication timing program. Here, using *Xenopus* egg extracts, we showed that Yap (Yes-associated protein 1), a downstream effector of the Hippo signaling pathway, is required for the control of DNA replication dynamics. We found that Yap is recruited to chromatin at the start of DNA replication and that Yap depletion accelerates DNA replication dynamics by increasing the number of activated replication origins. Furthermore, we identified Rif1, a major regulator of the DNA replication timing program, as a novel Yap binding protein. In *Xenopus* embryos, using a Trim-Away approach during cleavage stages devoid of transcription, we found that both Yap and Rif1 depletion trigger an acceleration of cell divisions, suggesting a shorter S-phase by alterations of the replication program. Finally, our data show that Rif1 knockdown leads to defects in the partitioning of early versus late replication foci in retinal stem cells, as we previously showed for Yap. Altogether, our findings unveil a non-transcriptional role for Yap in regulating replication dynamics. We propose that Yap and Rif1 function as breaks to control the DNA replication program in early embryos and post-embryonic stem cells.

**Highlights:**

- Yap is recruited to chromatin during DNA replication dependent on pre-replicative complex assembly.
- Yap controls DNA replication dynamics by limiting origin firing.
- The replication timing regulatory factor 1, Rif1, is a novel Yap binding-partner.
- Both Yap and Rif1 regulate the length of the first embryonic cell cycles.
- Like Yap, Rif1 controls retinal stem cell DNA replication timing.

## Introduction

Prior to cell division, DNA must be entirely and accurately duplicated to be transmitted to the daughter cells (Fragkos et al., 2015). In metazoan cells, DNA replication initiates at several thousands of fairly specific sites called replication origins in a highly-orchestrated manner in time and space (Machida et al., 2005; Prioleau and MacAlpine, 2016). In late mitosis and G1 phase, origins are first “licensed” for replication by loading onto chromatin the six ORC (origin recognition complex) subunits, then Cdc6 (cell-division-cycle 6) and Cdt1 (chromatin licensing and DNA replication factor 1), and finally the MCM (mini-chromosome maintenance) 2-7 helicase complex, forming the pre-replicative complex (pre-RC, for review see (Bell and Kaguni, 2013)). Pre-RC is subsequently activated during S-phase by cyclin- and Dbf4/Drf1- dependent kinases (CDKs and DDKs) which leads to the recruitment of many other factors, DNA unwinding and start of DNA synthesis at origins. In eukaryotes, segments of chromosomes replicate in a timely organized manner throughout S-phase. It is now widely accepted that the genome is partitioned into different types of genomic regions of coordinated activation (Marchal et al., 2019). During the first half of S-phase, the early-replicating chromatin, mainly transcriptionally active and localized to central regions of the nucleus, duplicates while late replicating chromatin, spatially located at the periphery of the nucleus, awaits until the second half (Berezney et al., 2000; Hiratani et al., 2010; Ryba et al., 2010). This pattern of DNA replication, also called DNA replication timing (RT) program, has been found to be stable, somatically heritable, cell-type specific, and associated to a cellular phenotype. As such, the RT can be considered as an epigenetic mark (Hiratani and Gilbert, 2009) and provides a specific cell state signature. Interestingly, a defined RT has been observed at very early stages in development, prior to the mid-blastula transition (MBT), in embryonic cells (also called blastomeres) undergoing rapid cell cycle consisting of only S/M phases that are typical of animals with external development (Siefert et al., 2017). Due to the absence of most transcriptional activities in these early embryos (Newport and Kirschner, 1982), the very rapid DNA synthesis during these cleavage divisions relies only on a stockpile of maternally supplied determinants. To date, little is known about the molecular cues that ensure faithful and complete DNA replication during early embryonic cell divisions. Very few gene knockouts have been shown to trigger alterations in the RT (Dileep et al., 2015; Marchal et al., 2019). Until now, the replication timing regulatory factor 1, Rif1, is one of the very few trans-acting factors whose loss of function has been found to result in major RT modifications in multicellular organisms (Cornacchia et al., 2012; Yamazaki et al., 2012). Rif1 inhibits the firing of late origins by targeting PP1 (Protein Phosphatase 1) to those origins, counteracting Cdc7/Dbf4 dependent Mcm4 phosphorylation in budding yeast and *Xenopus* egg extracts (Alver et al., 2017; Davé et al., 2014; Hiraga et al., 2014). We previously identified Yap as another factor implicated in RT control (Cabochette et al., 2015). Yap is a downstream effector of the Hippo signaling pathway. It was initially identified as a primary regulator of organ growth due to its action on embryonic progenitor cells (Huang et al., 2005; Lian et al., 2010; Ramos and Camargo, 2012; Zhao et al., 2010). Yap is mostly known to exert its function as a transcriptional co-activator acting via binding to the TEADs (transcriptional enhanced associated domain transcription factors) to control transcriptional programs involved in cell proliferation, differentiation, survival and migration (Totaro et al., 2018; Zhao et al., 2008). We previously found that *yap* is specifically expressed in neural stem cells in the *Xenopus* retina and that its knockdown in these cells leads to altered RT associated with a dramatic S-phase shortening (Cabochette et al., 2015). However, whether Yap is directly involved in DNA replication dynamics and whether it could regulate DNA replication during early embryonic divisions in the absence of transcription remained to be investigated.

We first addressed this question by taking advantage of *Xenopus* egg extracts, a cell-free system that faithfully recapitulates all steps of DNA replication (Blow and Laskey, 2016, 1986). We and others previously found that in this system activated replication origins are spaced 5 to 15 kb apart and clustered in early- and late-firing groups of origins (Blow et al., 2001; Herrick et al., 2000; Marheineke and Hyrien, 2004). Here, we found that Yap is recruited onto chromatin at the onset of DNA replication in *Xenopus* egg extracts in a manner that is dependent on the pre-RC formation. Yap depletion altered replication dynamics by increasing the number of activated origins. We further showed that Yap and Rif1 co-immunoprecipitated. As previously shown *in vivo* for *yap* (Cabochette et al., 2015), we found that *rif1* is expressed in retinal stem and early progenitor cells and involved in their RT signature. Finally, targeted protein depletion at early stages of embryonic development using a Trim-Away strategy, revealed the crucial role of both Yap and Rif1 in controlling the speed of cell divisions before MBT *in vivo*. Altogether, our findings unveiled Yap implication in the regulation of replication origin activation and identified Rif1 as a novel partner. We propose that Yap, like Rif1, acts as a brake during replication, to control the overall rate of DNA synthesis in early embryos and post-embryonic retinal stem cells.

## Results

### Yap is recruited to chromatin in a pre-RC-dependent manner in *Xenopus* egg extracts

Since Yap was described as a co-transcriptional factor, we wondered whether Yap was present in *Xenopus* egg extracts that are almost devoid of transcriptional activity (Wang and Shechter, 2016). By quantitative western blot, we found that Yap protein is present in S-phase egg extracts at a concentration of 11 ng/µl (169 nM, Figure 1 - figure supplement 1). We therefore further investigated the role of Yap during S-phase in this well characterized *in vitro* replication system where upon addition of sperm DNA to egg extracts, chromatin is assembled, replication proteins are imported, recruited on chromatin and nuclei synchronously start DNA replication. Thus, this *in vitro* system mimics the first embryonic S-phase. To know whether Yap interacts with chromatin during S-phase, we incubated sperm nuclei in egg extracts and collected purified chromatin fractions starting from pre-RC assembly up to ongoing DNA replication. This analysis revealed that Yap recruitment onto chromatin coincided with the loading of PCNA, an indicator of the recruitment of DNA polymerases and the start of DNA synthesis (Figure 1A). Yap further accumulated on chromatin following the progression of S-phase. Our results also showed that Yap is recruited to chromatin after the recruitment of the MCM complex (Mcm2, Mcm7) in the *Xenopus in vitro* system (Figure 1A). To address whether the recruitment of Yap could be dependent on pre-RC assembly on chromatin, we added to the egg extracts recombinant geminin, an inhibitor of Cdt1 necessary for MCM loading (McGarry and Kirschner, 1998; Tada et al., 2001). As a result, the recruitment of Yap on chromatin was severely delayed (Figure 1A, B). Thus, Yap is recruited to chromatin at the start of DNA replication and its recruitment is dependent on functional pre-RC assembly in the *Xenopus* egg extract system.

**Figure 1.**
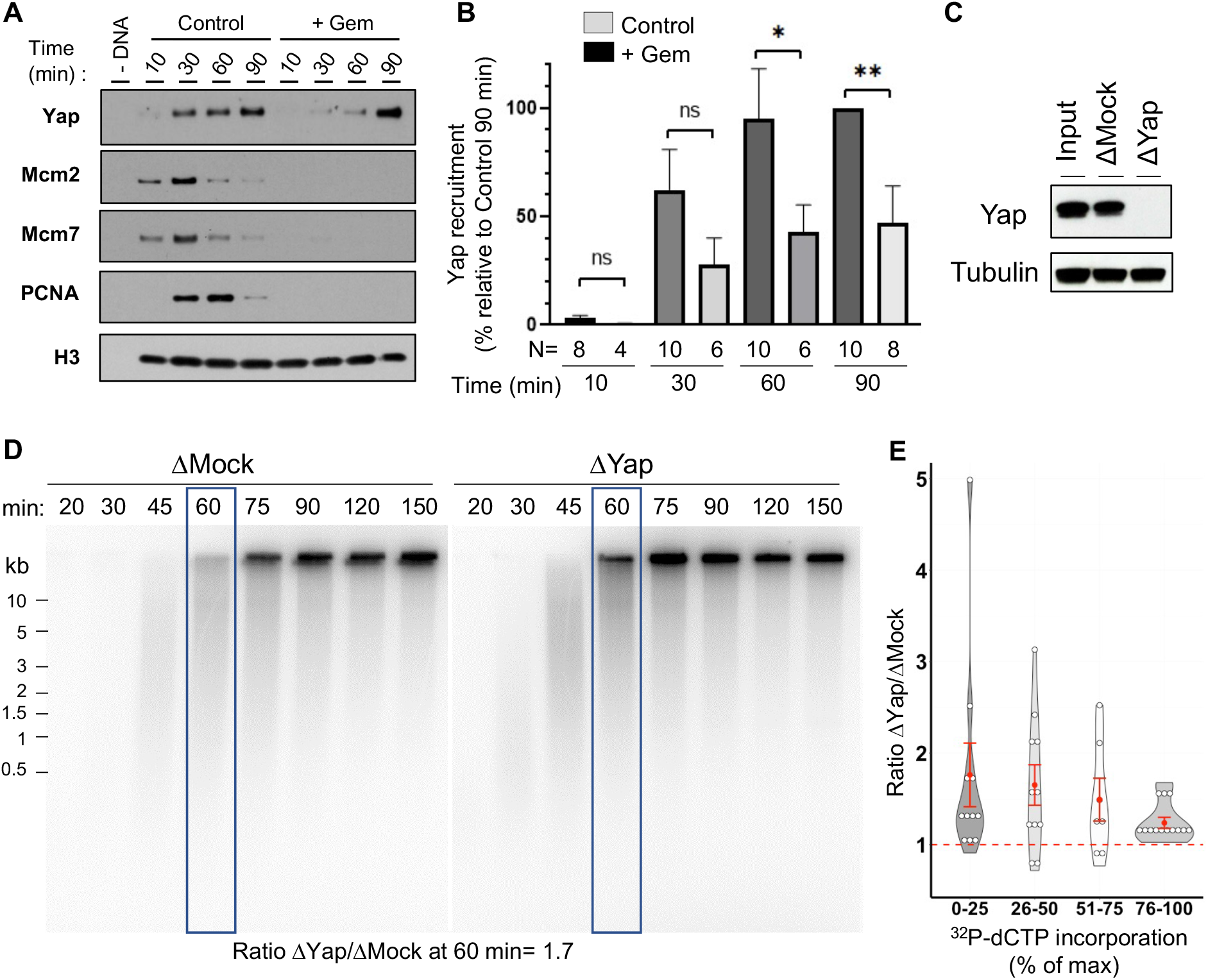
Yap is recruited to chromatin during DNA replication and the absence of Yap accelerates DNA synthesis in *Xenopus* egg extracts. (**A**) Sperm nuclei were incubated in *Xenopus* egg extracts in the absence (Control) or presence of geminin (+ Gem). Chromatin was isolated for immunoblotting at indicated times points before and during DNA replication. (**B**) Quantification of the amount of Yap: percentage of quantified optical densities of the Yap bands relative to that in the control condition at 90 minutes in isolated chromatin fractions. The number of analyzed fractions per time point during DNA replication is indicated in each bar (N=). ** p=0.0013; * p=0.0225; ns, not significant. Data is reported as mean ± SEM. (**C**) Western blot showing the efficiency of Yap protein depletion in *Xenopus* egg extracts. Extracts were immunodepleted with either a rabbit anti-Yap antibody (ΔYap) or a random rabbit IgG as a control (ΔMock). Tubulin is used as a loading control. (**D**) Immunodepleted extracts were supplemented with sperm nuclei and incubated with [α^32^P]dCTP for different times in order to label nascent DNA during replication. Nascent DNA strands synthesized were analyzed by alkaline gel electrophoresis. The level of radioactivity incorporation was quantified as % of maximal incorporation for each lane and the ratio of these values in ΔYap over ΔMock conditions was calculated for each time point. The ratio at 60 minutes is indicated as an example. **(E)** Violin plot showing ΔYap/ΔMock ratios from 8 independent experiments including the one depicted in (D). The time scale was fractionated in 4 periods to distinguish early, mid, late and very late phases of the replication process. The red dashed line highlights a ΔYap/ΔMock ratio of 1 that indicates no difference in the level of DNA synthesis between the 2 conditions, with the red dot indicating the mean and the red error bar the SEM, Wilcoxon signed ranked test, p-values: p= 0.002 (0-25%, n=10), p=0.014 (26-50%, n=11), p=0.16 (51-75%, n=6), p=0.0002 (76-100%, n=13).

### Yap depletion triggers the acceleration of DNA synthesis in egg extracts

To directly assess the role of Yap in DNA replication, we performed immunodepletion experiments. We were able to efficiently remove Yap from egg extracts (Figure 1C). We then used those Yap-depleted (ΔYap) or mock-depleted (ΔMock) egg extracts to monitor nascent strand DNA synthesis after incubating sperm nuclei in the presence of ^32^P-dCTP (Figure 1 D). Replication reactions were stopped at indicated times during S-phase and quantified (Figure 1E). We found that Yap depletion increased DNA synthesis during the early stages of DNA replication (30-60 min: low molecular weight nascent strands) and to a lesser extent at later stages (75-150 min: high molecular weight strands). We calculated the ratio between Yap- and Mock-depleted maximal incorporation at four different intervals of percentages of incorporation, reflecting early (0-25 % max. incorporation), mid (26-50 %), late (51-75 %) and very late (76-100%) S-phase. We found that Yap depletion increased on average DNA synthesis 1.8-fold during early S-phase, 1.7-fold during mid S-phase, 1.6-fold during late S-phase and 1.2-fold during very late S-phase. The increase in DNA replication after Yap depletion could be due to a quicker entry into S-phase, because of a more rapid chromatin assembly, rather than an effect on DNA replication itself. We however ruled out this hypothesis by analyzing nascent strands during very early S-phase, which did not reveal any precocious start of DNA synthesis after Yap depletion (Figure 1 – figure supplement 2). Of note, the effect of the Yap depletion was not rescued by adding back the recombinant protein (Figure 1 – figure supplement 3). Since Yap localization and function can be modified by many different post-translational modifications (PTMs) (Yan et al., 2020), one cannot exclude that one or more PTMs of the recombinant Yap produced in baculovirus-infected insect cells were missing. Altogether, we found that Yap depletion leads to accelerated DNA synthesis, mainly during the early to mid-stages of S-phase, suggesting that Yap negatively regulates the progression of DNA replication.

### Yap depletion increases replication origin firing

The higher rate of DNA synthesis observed in the absence of Yap could result from either an increase in origin firing, an increase in fork speed, or both. To directly monitor origin activation on single DNA molecules, we performed DNA combing experiments in control and Yap depleted extracts and determined the replication content, fork density, distances between replication eyes and eye lengths (Figure 2A, B, Figure 2 - table supplement 1). After Yap depletion, the mean replicated fraction significantly increased during early and mid S-phase by 2.5-fold (Figure 2B, C), consistent with the nascent strand analysis shown in Figure 1C. Moreover, Yap depletion significantly increased the mean density of active replication forks (1.8-fold; Figure 2D), demonstrating that the absence of Yap leads to an increase of activated replication origins. In parallel, a significant decrease in eye-to-eye distances occurred (ETED; Figure 2E). The increase of the overall fork density was more pronounced than the decrease of distances between neighbor origins analyzed at all time points (Figure 2, Figure 2 - table supplement 1). Therefore, this observation pointed to a role of Yap in regulating the activation of origins in not yet active, later replicating groups of origins. Replication eye lengths (EL) were not significantly different after Yap depletion at very early S-phase (Figure 2F), suggesting that fork speed was unchanged. Larger eye sizes detected at later time points (Figure 2 - table supplement 1) are most probably due to fusions of eyes from neighbor origins due to increased origin activation after Yap depletion, since we were not able to detect larger nascent strands in Yap depleted extracts during very early S-phase (Figure 1 – figure supplement 2). Altogether, we conclude that Yap depletion leads to an increase in origin activation, suggesting that Yap plays a key role in limiting origin firing during DNA replication.

**Figure 2.**
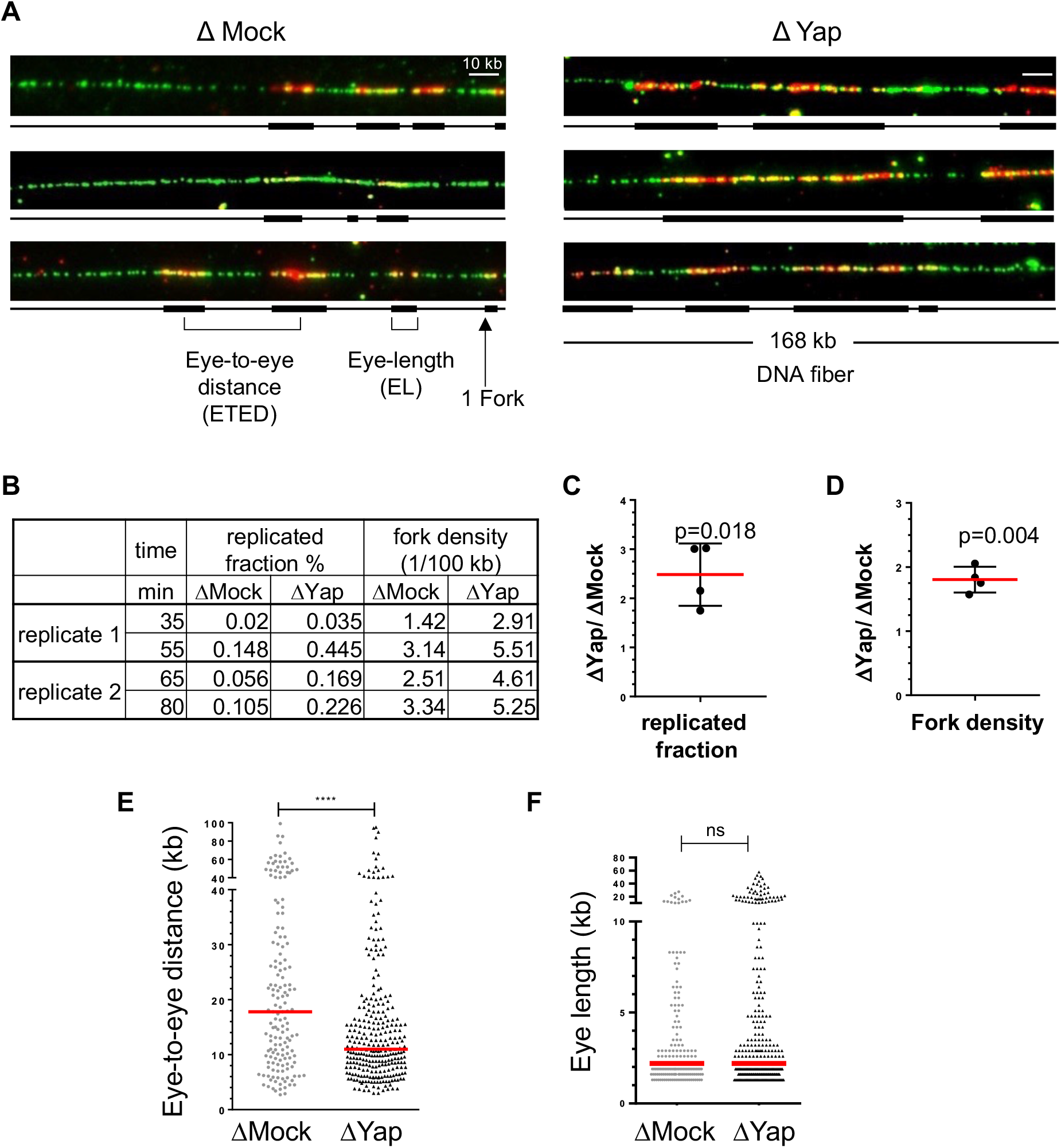
Egg extracts lacking Yap exhibit more active replication origins. Sperm nuclei were incubated in egg extracts in the presence of Biotin-dUTP and DNA combing was performed. **(A)** Three representative combed DNA fibers replicated in either the ΔMock- or ΔYap-depleted extracts (green: whole DNA labelling; red: biotin labelled replication eyes). **(B)** Replicated fraction and fork density (1/100 kb) of two independent experiments at 2 time points were calculated and reported in the table. **(C, D)** Corresponding scatter blots of ΔYap/ΔMock ratios of replicated fraction (C) and fork density (D), with mean (n=4) and standard deviation, P values, one sample t-test, compared to theoretical mean 1. **(E, F)** The eye-to-eye distance distributions after mock or Yap depletion (E, ETED, scatter dot plots with median, n=157 or 286, replicate 2, 80 min) and the eye length distributions (F, EL, scatter dot plots with median, n=182 or 311, replicate 2, 65 min).

### Yap interacts with Rif1

To identify Yap partners in the context of DNA replication, we conducted an exploratory search for Yap-interacting proteins by co-immunoprecipitation coupled to mass spectroscopy (co-IP-MS) in S-phase egg extracts (see data availability). Among the proteins enriched more than 3-fold in Yap-co-IP versus control-co-IP conditions, we mostly identified factors functionally associated with mRNA metabolic process, ribonucleoprotein complex assembly and translation (Figure 3A). This is in accordance with the fact that *Xenopus* egg extracts possess little or no intrinsic transcriptional activity but can strongly support translation and post-translational modification (Matthews and Colman, 1991). Of note, our analysis did not point to GO term enrichments related to DNA replication *per se*. However, we identified an interesting candidate, Rif1, a major factor of the replication timing program (Cornacchia et al., 2012; Yamazaki et al., 2012). Interestingly, both Yap and Rif1 are associated with the stem cell population maintenance GO term.

**Figure 3.**
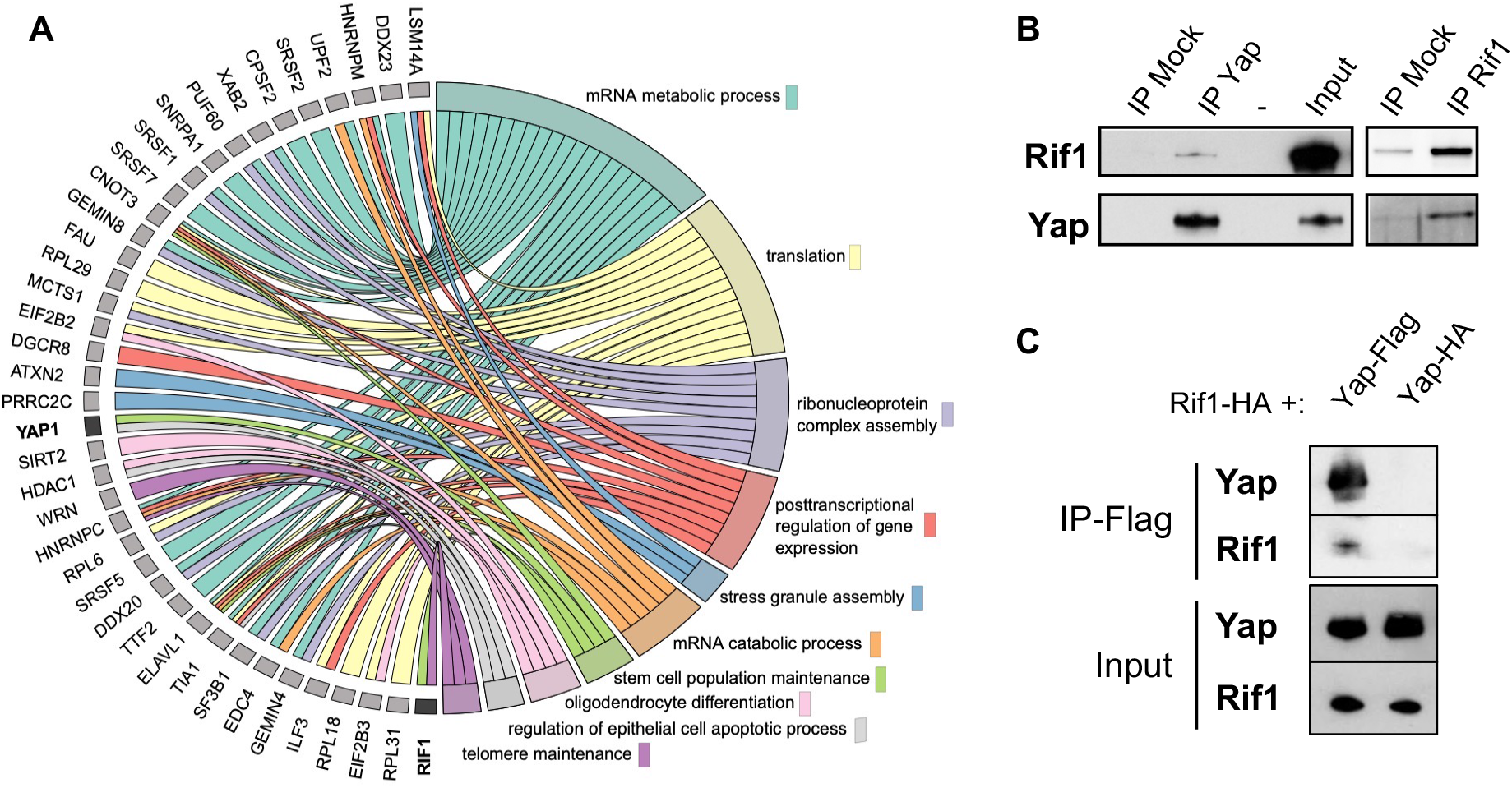
Rif1 interacts with Yap. (**A**) Chord plot representation related to GO annotations belonging to biological processes of proteins enriched by at least 3-fold in Yap versus control co-immunoprecipitations performed in S-phase egg extracts. Note that Yap and Rif1 are both functionally associated with stem cell population maintenance. (**B**) Anti-Yap (IP Yap), anti-Rif1 (IP Rif1) or control (IP Mock) antibodies coupled to Sepharose beads were incubated in S-phase egg extracts; immunoprecipitates were subjected to gel electrophoresis and western blotted using the indicated antibodies. -, unloaded lane. (**C**) Extracts from HEK293T cells transfected with the indicated tagged constructs were immunoprecipitated using anti-Flag antibody. The input and immunoprecipitates were subjected to gel electrophoresis and western blotted using the indicated antibodies.

We confirmed this Yap/Rif1 interaction in egg extracts by reciprocal co-immunoprecipitation assays (Figure 3B). We next showed that this interaction between Rif1 and Yap also exists following the expression of the tagged proteins in HEK293 cells (Figure 3C). Altogether, our data uncovered Rif1 as a Yap interacting factor, supporting the role of Yap in the regulation of DNA replication dynamics.

### Yap and Rif1 depletions accelerate cell division rate *in vivo* in embryonic cells

To assess whether Yap non-transcriptional function in DNA replication dynamics also holds true *in vivo*, we took advantage of the early cell divisions of *Xenopus* embryos that provide a simplified system of the cell cycle. Indeed, during early development prior to the mid-blastula transition (MBT, stage 8), cells divide very rapidly, rather synchronously for a series of 12 divisions and present a cell cycle structure without gap phases. As a result, variations of the number of cells at a given time during this developmental period reflect alteration of the time spent in the S and M phases. We thus decided to deplete embryos from Yap and assess the outcomes on the rate of embryonic cell division. Since Yap protein is maternally expressed (Figure 4A), we employed the recently developed Trim-Away technique (Clift et al., 2018, 2017; Weir et al., 2021) to directly trigger *in vivo* the degradation of Yap protein stockpile (Figure 4A). The Trim-Away mediated knockdown has previously been shown to be effective in *Xenopus* for another target using Trim21 mRNAs (Weir et al., 2021). Here, we decided to use the Trim21 protein instead, to prevent delay inherent to the translation process. In addition, since we observed an increase in the level of Yap protein before MBT (Figure 4A), we combined the Trim-Away approach with injections of *yap* translation blocking morpholino oligonucleotides (MO) to prevent *de novo* protein synthesis (Figure 4B). We found that Yap degradation was effective from the 8-cell stage onwards using the Trim-Away approach and that the combined strategy (Trim-Away + MO) led to a stronger and prolonged Yap depletion (Figure 4 – figure supplement 1A). We then monitored the progression of cell division before MBT. We found that cells were smaller and more numerous in Yap depleted embryos than in controls at stage 7 (Figure 4C, D). This suggests that Yap depletion leads to an increased speed of cell divisions in pre-MBT *Xenopus* embryos. The resulting embryos do develop but they display severe abnormalities by the tadpole stage (Figure 4 – figure supplement 1B).

**Figure 4.**
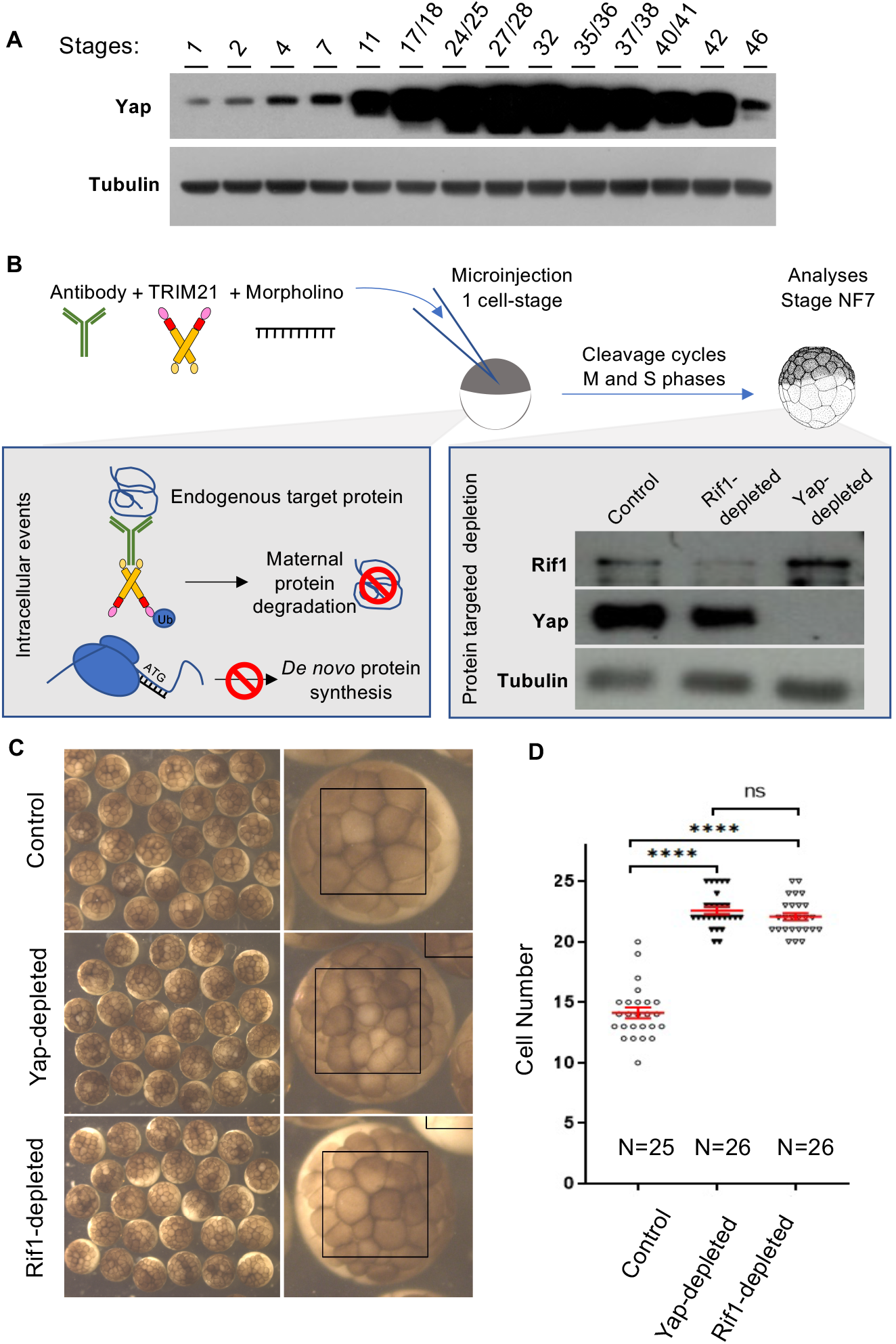
Yap and Rif1 depletion accelerate cell cycles in early *Xenopus* embryos. (**A**) Time course analysis of Yap expression throughout development by western blot. (**B**) Diagram of the experimental procedure used in (C) and western blot showing the efficiency of *in vivo* Yap and Rif1 depletions at stage 7 using combined Trim-Away and MO-mediated knock-down approaches. *X. laevis* embryos were microinjected at the one-cell stage with (i) control MO + pre-immune serum + TRIM21 (Control), (ii) *Yap*-MO + anti-Yap antibody + TRIM21 (Yap-depleted), or (iii) *rif1*-MO + anti-Rif1 antibody + TRIM21 (Rif1-depleted). (**C**) Images from representative injected stage 7 embryos. A close-up view is shown on the right panels for each condition. (**D**) The number of cells per embryo within the area defined in B (black boxes) was quantified. Data are shown as individual value plots with error bars (mean with SEM in red). Mann-Whitney test; **** p ≤ 0.0001; ns, not significant Scale bar = 500 µm.

We wondered whether the depletion of Rif1 could lead to a similar phenotype. Rif1 depletion was previously shown to increase the rate of DNA replication in *Xenopus* egg extracts during unchallenged S-phase (Alver et al., 2017), but it was unknown how Rif1 depletion would affect early embryonic cell cycles *in vivo*. We undertook the same strategy to deplete Rif1 from *Xenopus* embryos using both the Trim-Away technique and *rif1*-MO. We found that Rif1 depletion from embryos also led to an increased number of cells at stage 7, indicative of a faster rate of cell division, as previously observed upon Yap depletion (Figure 4 C, D). Considering the known function of Rif1 in DNA replication and the absence of gap phases in pre-MBT embryos, our results strongly suggest that the increased rate of cell division in absence of Rif1 results from an acceleration of DNA replication and a shortening of S-phase length. We therefore propose that both Yap and Rif1 are involved in controlling the DNA replication dynamics in pre-MBT embryos.

### *rif1* is expressed in retinal stem cells and its knockdown affects their temporal program of DNA replication

Since we found this new interaction between Rif1 and Yap and since Rif1 has been recently shown to function in a tissue-specific manner (Armstrong et al., 2020), we investigated its expression and function in *Xenopus* retina and compared the results with our previous findings regarding Yap retinal expression/function (Cabochette et al., 2015). *In situ* hybridization study and immunostaining experiments revealed prominent *rif1* expression in the periphery of the ciliary marginal zone (CMZ) of the retina (Figure 5A-C), a region containing stem and early progenitor cells (Perron et al., 1998), and where *yap* is also specifically expressed (Cabochette et al., 2015). We next undertook a knockdown approach using *rif1*-MO (Figure 5D-F). Morphant tadpoles exhibited significantly reduced eye size compared to controls (Figure 5E, F), as did *Yap* morphants (Cabochette et al., 2015). Importantly, in support of the specificity of *rif1*-MO, this phenotype was restored upon co-injection of a *rif1*-MO with non-targetable *rif1* mRNAs (Figure 5 – figure supplement 1).

**Figure 5.**
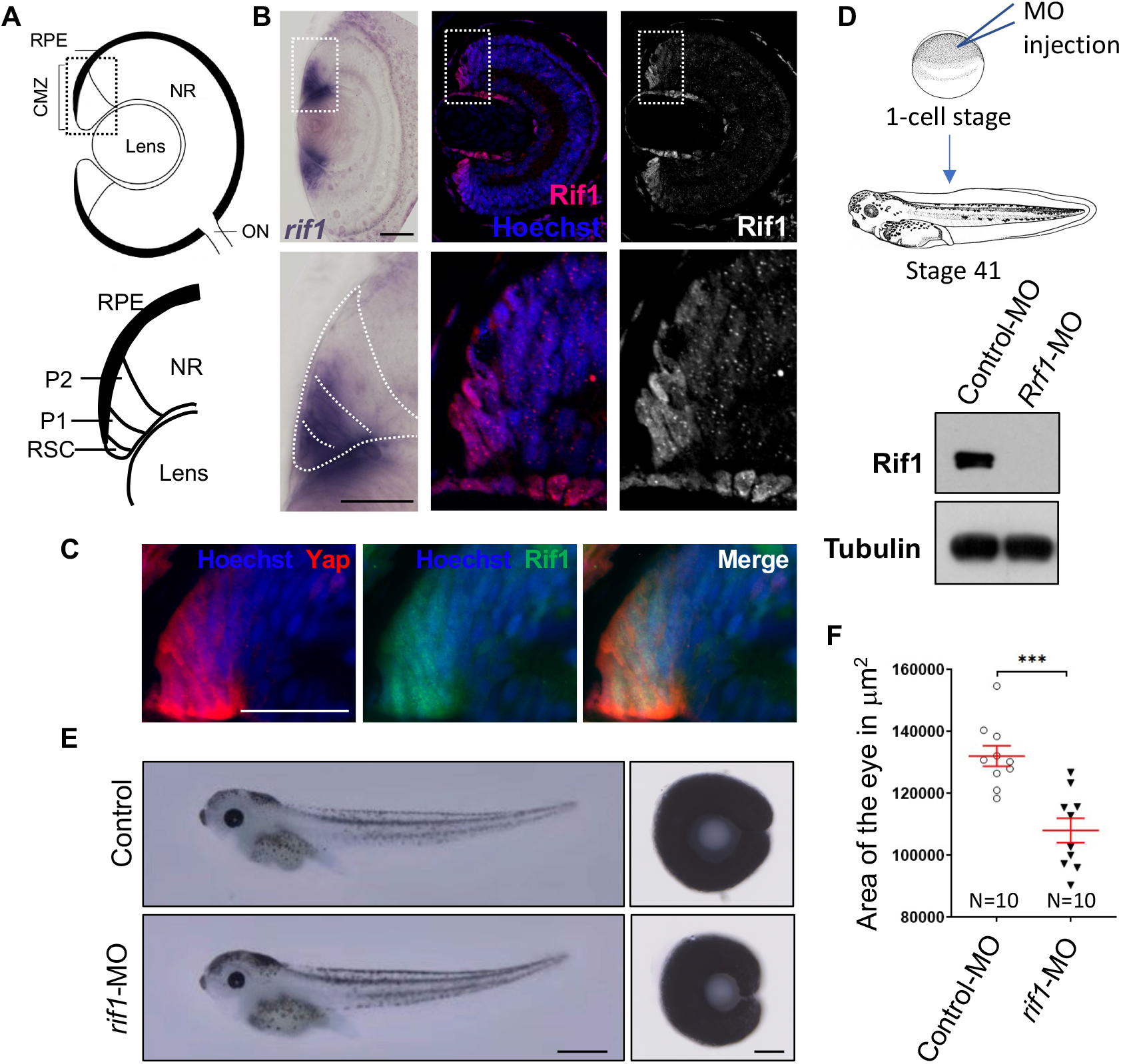
*rif1* is expressed in retinal stem cells and its knock-down leads to small eye phenotype. **(A)** Schematic transversal section of a *Xenopus* tadpole retina (RPE: retinal pigmented epithelium; NR: neural retina; ON: optic nerve). Within the ciliary marginal zone (CMZ; lower diagram), retinal stem cells (RSC) reside in the most peripheral margin while early (P1) and late (P2) progenitors are located more centrally. (**B**) Retinal sections from stage 41 *Xenopus* tadpoles following *in situ* hybridization for *rif1* expression (left panels, in purple) or immunostained for Rif1 (middle panel in red along with nuclei counterstained with Hoechst in blue, and right panel Rif1 alone in white). The images on the lower panels are higher magnification of the CMZ (delineated dotted lines on the top panels). (**C**) CMZ region of retinal sections from stage 41 *Xenopus* tadpoles co-immunostained for Yap (red) and Rif1 (green) along with nuclei counterstained with Hoechst (blue). (**D**) Diagram showing the experimental procedure used in (E). One cell-stage embryos are microinjected with Control MO or *rif1*-MO and analyzed at stage 41. The western blot shows the efficiency of the MO at depleting Rif1 in embryos. (**E**) Tadpoles microinjected with MO as shown in (D) and corresponding dissected eyes (right panels). (**F**) The area of dissected eyes was measured for 10 embryos per condition. Data are shown as individual value plots with error bars (mean with SEM in red). Mann-Whitney test; *** p=0.0002. Scale bar = 50 µm in (B, C), 1 mm in (E, tadpoles) and 100 µm in (E, dissected eyes).

We next analyzed the level of proliferation within the CMZ in *rif1* morphant tadpoles (Figure 6). Unlike the observed decreased of the EdU cell number in *yap* morphant CMZ (Cabochette et al., 2015), we did not find here any significant difference in the number of EdU+ cells in *rif1*-MO-injected tadpoles compared to controls (Figure 6E). Interestingly however, as observed in *yap* morphants (Cabochette et al., 2015), we found a drastic change in the distribution of EdU-labelled replication foci in retinal stem and early progenitor cells, where *rif1* is normally expressed (Figure 6 A-D, F). Short pulse labelling experiments indeed allow the visualization of replication foci in cells. The spatial distribution of these foci evolves in a stereotyped manner during S-phase (Figure 6A): from numerous small foci located throughout the nucleus in early S-phase, to few large punctuated ones in mid/late S-phase (Koberna et al., 2005; Van Dierendonck et al., 1989). Our analysis revealed a decreased proportion of cells exhibiting a mid-late versus early S-phase patterns in *rif1* morphants compared to controls (Figure 6F). We thus propose that, like *yap* knockdown, *rif1* knockdown alters the spatial organization of replication foci in CMZ cells, suggesting that both Yap and Rif1 may control the RT program *in vivo* in post-embryonic retinal stem/early progenitor cells.

**Figure 6.**
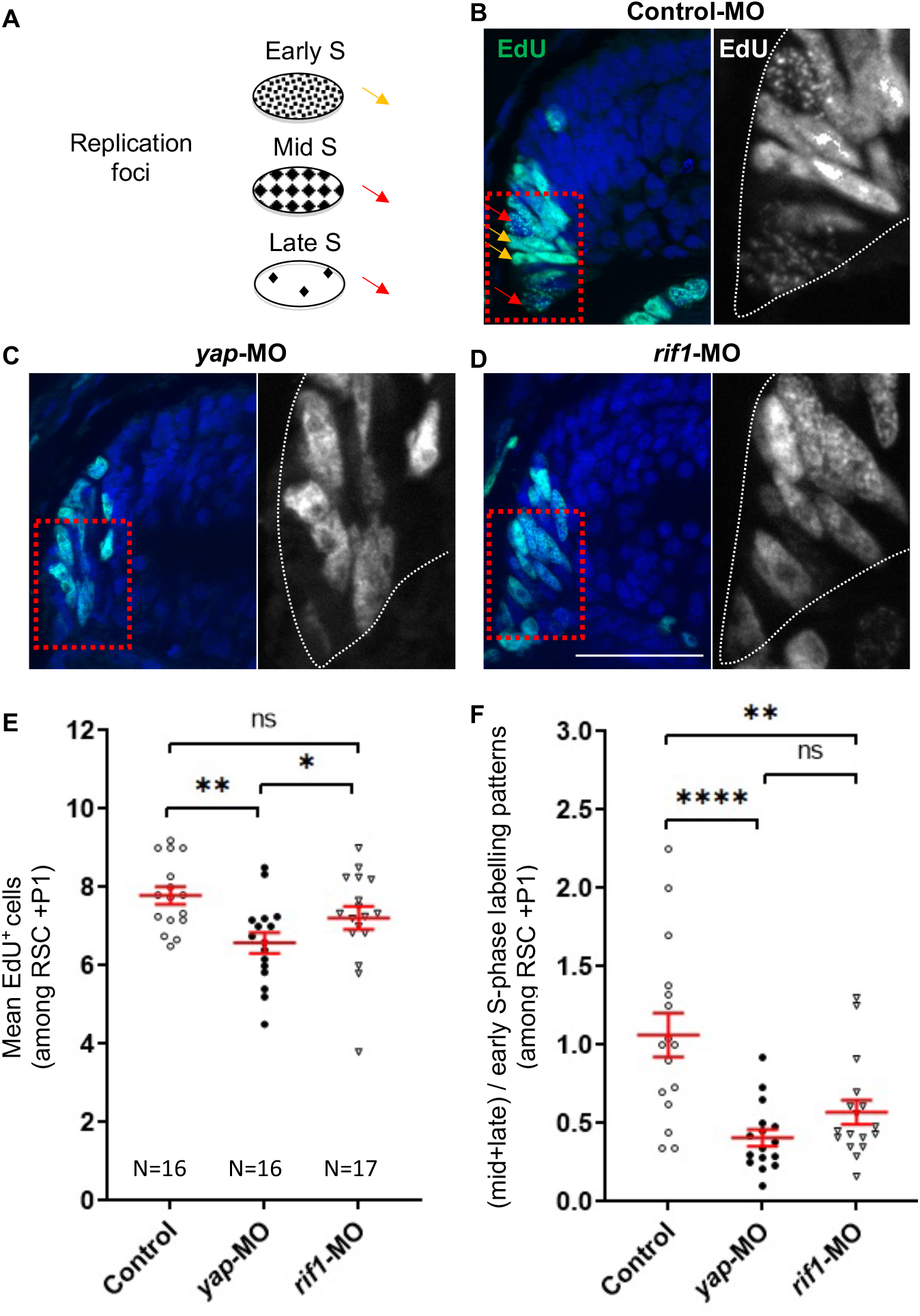
*rif1* loss of function affects DNA replication timing in retinal stem/early progenitor cells. **(A)** Schematic representation of the replication foci observed during S-phase progression as inferred from EdU labelling. Orange arrows indicate typical early S replication patterns and red arrows, mid/late S replication ones (only 2 examples of each are shown in panel B). (**B-D**) One cell-stage embryos are microinjected with either the control MO (**B**), *yap*-MO (**C**) or *rif1*-MO (**D**) and analyzed for EdU-labelled replication foci (1 hr.-pulse) at stage 41. The region enlarged on right panels is delineated with red dashed lined boxes. The tip of the CMZ is highlighted by dotted white lines in the enlargements. (**E, F**) Quantifications of EdU^+^ cell number (E) and of the ratio of mid+late/early-like foci patterns (F). In both cases, only cells in the RSC and P1 regions (see diagram shown in Figure 5A) have been analyzed. The number of analyzed retinas is indicated for each condition. Data are shown as individual value plots with error bars (mean with SEM in red). Mann-Whitney test; ** p≤0.01, * p=0.05; in **** p≤0.0001; ns, non-significant. Scale bar = 50 µm.

## Discussion

During S-phase, eukaryotic DNA is not replicated all at once, but large genomic regions are duplicated in a characteristic temporal order known as the RT program. To date, very few factors involved in the orchestration of this program have been identified. We have previously revealed that *yap* knockdown is associated with an altered RT program in *Xenopus* retinal stem cells (Cabochette et al., 2015). Whether and how Yap could directly regulate DNA replication was however unknown. Here, we used the *Xenopus in vitro* replication system and early *Xenopus* embryos, where DNA transcription is absent, to study the role of Yap in S-phase, independently from its transcriptional function. Our study shows that Yap regulates DNA replication dynamics in these embryonic systems. First, we found that Yap is recruited to chromatin at the start of DNA synthesis, and this is dependent on the pre-replicative complex assembly. Second, our data revealed a non-transcriptional role for Yap in the initiation of DNA replication. Third, we identified Rif1, a global regulator of the RT program, as a novel Yap partner. Finally, our *in vivo* data suggest that Yap and Rif1 are similarly involved in both the spatial organization of DNA replication foci and DNA replication dynamics. We propose a model in which Yap and Rif1 would limit replication origin firing and as such act as breaks during S-phase to control the overall rate of DNA synthesis (Figure 7).

**Figure 7.**
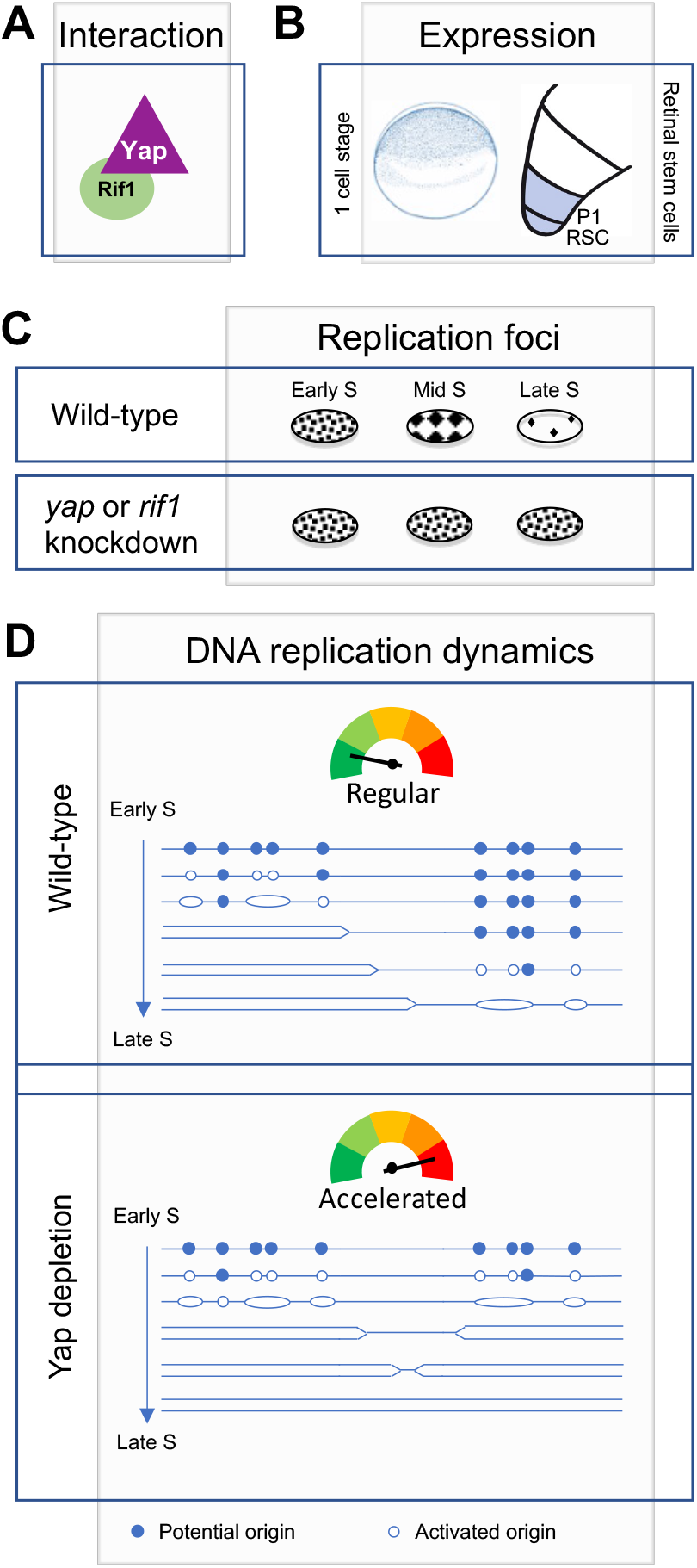
Diagram illustrating Yap function in the control of DNA replication dynamics. We found that Yap and Rif1 can interact (**A**) and are co-expressed in *Xenopus* early embryos as well as in retinal stem/progenitor cells (**B**). (**C**) We found that *yap* and *rif1* knockdowns in retinal stem/progenitor cells similarly alter the proper repartition of early and late-like patterns of replication foci (this study and (Cabochette et al., 2015)). (**D**) We propose a model where Yap would ensure the proper orchestration of the RT program during early development. The schematic representation of the replication program was adapted from Gaboriaud J. and Wu PJ (Gaboriaud and Wu, 2019). Based on our assays *in vitro* in egg extracts and *in vivo* in early embryos, we propose that following Yap depletion (bottom panel), the number of firing origins is increased and S-phase length is reduced compared to a wild-type situation (top panel).

The molecular control of the RT program remains elusive. Regarding key factors, Rif1 was the first mammalian factor shown to temporally control DNA replication, acting negatively on origin activation (Cornacchia et al., 2012; Hayano et al., 2012; Yamazaki et al., 2013, 2012). This function of Rif1 depends on its interaction with protein phosphatase 1 (PP1) (Davé et al., 2014) which counteracts the DDK dependent activation of MCM2-7. On the other hand, Polo-like kinase 1 (Plk1) positively controls replication origin firing by negatively regulating Rif1-PP1 interaction (Ciardo et al., 2021b). Here, we found that Yap is a novel key component of this molecular machinery that regulates replication origin activation. First, we found that Yap and Rif1 interact physically. Whether this interaction could impact Rif1-PP1 association remains to be determined. Interestingly, it has previously been shown that PP1 interacts with and dephosphorylates Yap (Wang et al., 2011), suggesting the potential existence of a Yap-Rif1-PP1 multi-protein complex. Second, we found that Yap depletion leads to similar DNA replication dynamics defects than those obtained following Rif1 depletion in early embryos. Third, in egg extracts Yap depletion increases the overall rate of DNA replication, similarly to what was shown before in the absence of Rif1 in the same *in vitro* system (Alver et al., 2017). Finally, we also observed similar phenotypes following *rif1* or *Yap* knockdown in *Xenopus* early embryos (i.e. increase in cell cycle speed) and in retinal stem cells (i.e. increase in early-like replication foci patterns). It is thus tempting to speculate that Rif1 and Yap act in concert to regulate replication dynamics.

We observed that Yap is recruited to replication competent chromatin at the start of S-phase and accumulates over S-phase, consistent with a direct role in DNA replication regulation. This dynamic behavior is similar to the observed increase of chromatin bound Rif1 (Kumar et al., 2012). We showed that Yap loading on chromatin depends on a functional pre-RC assembly or on DNA replication *per se*, since inhibition of pre-RC assembly also inhibits S-phase entry. We however do not know how Yap is recruited to chromatin in the first place since our proteomic analysis did not reveal a direct interaction with any members of the pre-RC complex. Therefore, Yap might be recruited by proteins involved in steps downstream of pre-RC assembly. We found that the increased rate in DNA synthesis after Yap depletion is due to an increase in replication origin activation, especially early in S-phase, strongly suggesting that Yap directly limits origin-firing. Whether it prevents late origin firing in early S-phase cells or whether it inhibits dormant origin firing around active replication forks remains to be investigated. However, the increase in early-like foci patterns at the expense of late-like ones in retinal stem cells observed upon *yap* (Cabochette et al., 2015) or *rif1* knockdown (this study) rather suggests an impact on the partition between early and late replication-firing. In Rif1-depleted Hela cells, the overall replication foci were similarly found to be extensively rearranged, with cells displaying predominantly early S-phase-like patterns (Yamazaki et al., 2012). In addition, our DNA combing analysis after Yap depletion demonstrated that the overall fork density was increased to a higher extent than local origin distances were decreased. This suggests that Yap controls origin firing more at the level of groups of origins or replication clusters than at the level of single origins, therefore regulating more the temporal control of origin activation.

In embryos, the increased speed of cell division during early embryonic cleavage cycles that we observed upon *yap* or *rif1* knockdown is consistent with a function of both factors in limiting the replication of late genomic regions leading to a slowing down of S-phase. In retinal stem cells however, Yap and Rif1 do not seem to function similarly on S-phase length. Indeed, although knockdown phenotypes were similar in terms of early-late foci ratio regulation, only *Yap* knockdown led to a significant change in the proportion of S-phase cells among stem and progenitor cells (i.e. number of EdU+ cells at the tip of the CMZ). How S-phase length is differentially regulated in both cases following altered RT program remains to be investigated. It is well known that Yap also transcriptionally regulates cell cycle genes, which may contribute to such different outcomes. Interestingly, among direct targets genes regulated by Yap, there are also essential factors involved in replication licensing, DNA synthesis and repair (e.g. *CDC6, GINS1, MCM3, MCM7, POLA2, POLE3, TOP2A* and *RAD18*; (Zanconato et al., 2015)). Yap thus likely impacts replication dynamics at both the transcriptional and non-transcriptional levels in retinal stem cells.

Combined observations point to a role for Rif1 in higher order chromatin architecture and its relationship with RT (Foti et al., 2016). Rif1 localizes to late-replicating sites of chromatin and acts as a remodeler of the three-dimensional (3D) genome organization and as such defines and restricts the interactions between replication-timing domains (Foti et al., 2016). Although the RT program can be established independently of the spatial distribution of replication foci, nuclear organization and RT are correlated and Rif1 is central in co-regulating both processes (Gnan et al., 2021). It would thus be interesting in the future to assess whether Yap function in DNA replication could be linked to a role as an organizer of nuclear architecture. Recent studies invoke liquid-liquid phase separation (LLPS) in the establishment of chromatin activity and the formation of chromatin compartments (Rippe, 2021). Interestingly, Yap has been described to form liquid like condensates in the nucleus (Cai et al., 2019). Whether higher level replication organization is impacted by LLPS is currently unknown, but several members of the pre-RC complex (Orc1, Cdc6, Cdt1) are also able to induce DNA dependent liquid-liquid phase separations *in vitro* (Parker et al., 2019).

Not much is known about signaling pathways regulating the RT or Rif1 activity. The ATM/53BP1 signaling pathway has been identified upstream of the RT and relays information onto Rif1 activity in response to DNA double strand breaks (Kumar and Cheok, 2014). Future work will be required to assess whether Yap activity in the context of DNA replication is regulated by the Hippo pathway. Interestingly, LATS1, another component of the Hippo pathway, has been involved in the ATR-mediated response to replication stress (Pefani et al., 2014). Several Hippo pathway components may thus regulate, independently or in concert, replication dynamics.

The role of Yap and Rif1 in the regulation of the RT program in early embryos opens new questions regarding the dynamics of RT changes during development. Although it was previously thought that the spatio-temporal replication program is not established until the MBT (Hyrien et al., 1995; Sasaki et al., 1999), it was also demonstrated that the oocyte-type of 5S RNA genes replicate later than the somatic-types of 5S RNA genes in *Xenopus* egg extracts (Wolffe, 1993). It was then shown that the RT program in *Xenopus in vitro* system is not completely random, with large chromosomal domains being replicated in a reproducible manner (Labit et al., 2008). Moreover, in zebrafish embryos, a genome-wide approach clearly established that a compressed, yet defined, RT program is evident before the MBT (Siefert et al., 2017). Further on, it was recently demonstrated that the replication program is regulated at the level of large domains by the replication checkpoint (Ciardo et al., 2021a) and by polo-like kinase 1 via the inhibition of Rif1/PP1 (Ciardo et al., 2021b). Altogether, those findings strongly suggest the existence of an embryonic RT program before the MBT. Remarkably, gradual changes in the RT program occur from the MBT and throughout development. The molecular control behind this dynamic is unknown. How Yap and Rif1 functions evolve at different stages and impact changes in the RT program at this important transition is therefore an interesting issue to be addressed in the future. To do so, the new combination of tools implemented in this study will be very valuable, as we have proven the efficient depletion of maternal proteins stockpiles while preventing their *de novo* synthesis by combining the Trim-Away technique with MO injections before MBT. In this context, *Xenopus* embryos seem particularly suitable to shed light and dissect the molecular mechanisms at work during embryogenesis that underlie RT changes. Identifying and characterizing the factors controlling these changes during development will certainly have an impact on how we approach questions related to cellular reprogramming, stem cells and cancer biology.

## Materials and Methods

### Ethics statement

All animal experiments have been carried out in accordance with the European Community Council Directive of 22 September 2010 (2010/63/EEC). All animal care and experimentation were conducted in accordance with institutional guidelines, under the institutional license C 91-471-102. The study protocols were approved by the institutional animal care committee CEEA #59 and received an authorization by the Direction Départementale de la Protection des Populations under the reference APAFIS#998-2015062510022908v2 for *Xenopus* experiments.

### Embryo, tadpole and eye collection

*Xenopus laevis* embryos were obtained by conventional methods of hormone-induced egg laying and *in vitro* fertilization (Sive et al., 2007), staged according to Nieuwkoop and Faber’s table of development (Nieuwkoop and Faber, 1994), and raised at 18-20°C. Before whole eye dissection, tadpoles were anesthetized in 0.005% benzocaine. Dissected eye area was measured using AxioVision REL 7.8 software (Zeiss).

### Antibodies and recombinant proteins

A detailed list of the antibodies used in this study for immunohistochemistry (IHC), immunodepletion and western blot (WB) is provided in Supplementary table. HLTV-hTRIM21 was a gift from Leo James (Addgene plasmid #104973; http://n2t.net/addgene:104973; RRID: Addgene_104973). Recombinant His-geminin, and His-hTRIM21 were prepared as described (respectively (Clift et al., 2017; Toyoshima and Hunter, 1994). C-terminal *Xenopus rif1* cloned in pET30a vector (a gift from W. Dunphy and A. Kumagai (Kumar et al., 2012)), was expressed in *Escherichia coli* C41 cells, purified by Nickel-Sepharose chromatography (Amersham Bioscience), and used as an antigen to raise antibodies in rabbits at a commercial facility (Covalab, Villeurbanne, France). A cDNA encoding recombinant His-tagged *Xenopus* Yap was cloned in pFastBac1vector, expressed in the baculovirus Bac-to-Bac expression system (Invitrogen), purified by Nickel-Sepharose chromatography as described by the supplier (Amersham Bioscience) and then dialyzed overnight against 25 mM Hepes pH 7.8, 250 mM NaCl, 5 mM imidazole, 5% glycerol, 7.5 mM MgCl2, 1 mM DTT, 1 mM EDTA. Purified His-Yap was then used as an antigen to raise antibodies in rabbits at a commercial facility (Covalab, Villeurbanne, France).

### Morpholinos and TRIM21 microinjections

For *in vivo* depletion experiments, 2 pmol of *Yap-*Morpholinos (MO, Gene Tools, LLC) or 1 pmol of *rif1*-MO or 2 pmol of standard control MO together with a fluorescent tracer (dextran fluorescein lysine, Thermo Fisher Scientific) were microinjected into fertilized eggs. The Trim-Away experiments were conducted in a similar way using a mixture of recombinant hTRIM21, anti-Rif1 or anti-Yap antibody together with 1 or 2 pmol of *rif1*-, *Yap-* or control-MO. MO sequences used in this study can be found in Supplementary table.

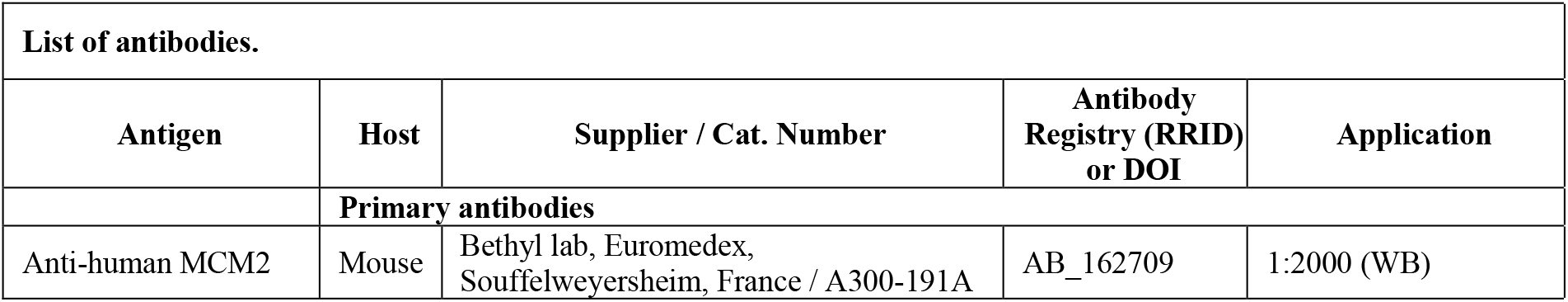
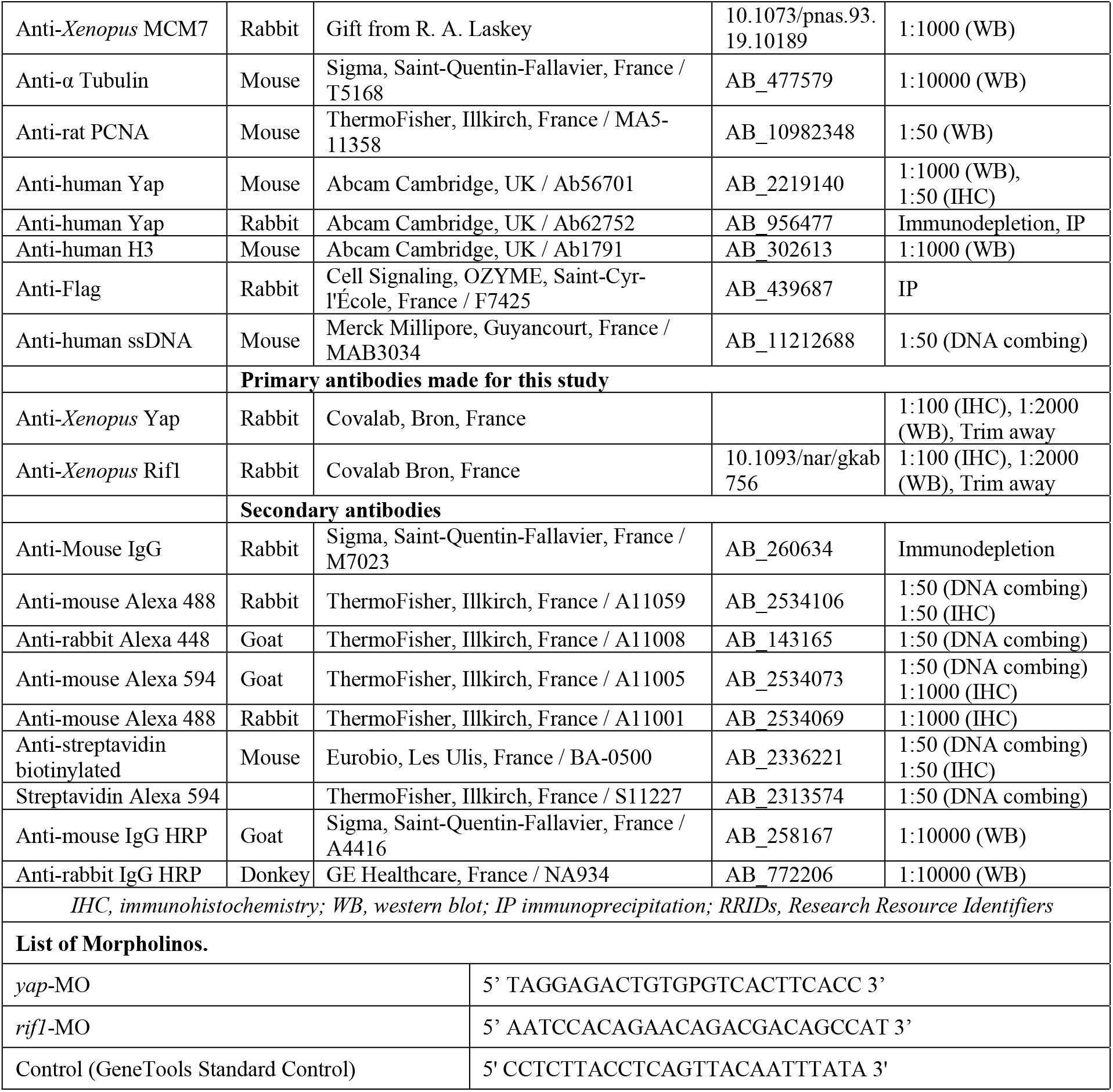

### Supplementary table Replication of sperm nuclei in *Xenopus* egg extracts

Replication competent extracts from unfertilized *Xenopus* eggs and sperm nuclei from testis of male frogs were prepared as described (Blow and Laskey, 1986). Sperm nuclei (2000 nuclei/µl) were incubated in untreated, mock or Yap depleted extracts in the presence of cycloheximide to inhibit translation (250 µg/ml, Sigma), energy mix (7.5 mM creatine phosphate, 1 mM ATP, 0.1 mM EGTA, pH 7.7, 1 mM MgCl2). Loading of the MCM complex (pre-RC assembly) was prevented by addition of 100 nM of recombinant geminin to the extracts.

### Immunodepletions

Rabbit anti-Yap antibody (ab62752, Abcam), pre-immune serum or rabbit IgG (M7023, Sigma) were coupled overnight at 4°C to protein A Sepharose beads (GE Healthcare). Coupled beads were washed three times in EB buffer (50 mM Hepes, pH 7.5, 50 mM KCl, 5 mM MgCl2) then incubated 1 hour at 4°C in egg extracts (volume ratio 1:3).

### Neutral and alkaline agarose gel electrophoresis

Sperm nuclei were incubated in fresh extracts complemented with indicated reagents and one-fiftieth volume of [α-^32^P]dCTP (3000 Ci/mmol). DNA was recovered after DNAzol® treatment (Invitrogen protocol) followed by ethanol precipitation, separated on 1.1% alkaline agarose gels, and analyzed as described (Marheineke and Hyrien, 2001). From one extract to another, the replication extent (percent of replication) differs at a specific time point, because each egg extract replicates nuclei with its own replication kinetics. In order to compare different independent experiments performed using different egg extracts, the data points of each sample were normalized to maximum incorporation value of 0-100 %. To include statistics, the scaled data points were grouped into 4 bins (0-25% = early; 26-50% = mid; 51-75% = late; 76-100% = very late S phase); mean and standard deviation were calculated for each bin and a Wilcoxon signed ranked test was used to assess statistically significant differences between the data in each bin.

### Western blot

For analysis of chromatin-bound proteins, we used a protocol slightly modified from (Räschle et al., 2008). Briefly, reactions were diluted into a 13-fold volume of ELB buffer (10 mM Hepes pH 7.5, 50 mM KCl, 2.5 mM MgCl2) containing 1 mM DTT, 0.2% Triton X100, protease inhibitors and phosphatase inhibitors. Chromatin was recovered through a 500 mM sucrose cushion in ELB buffer at 6780g for 50 sec at 4°C, washed twice with 200 µl of 250 mM sucrose in ELB buffer, and resuspended in 20 µl SDS sample buffer. Western blots were conducted using standard procedures on *Xenopus* embryo/tadpole protein extracts. Proteins were loaded, separated by 7.5%, 12% or 4-15% SDS-polyacrylamide gels (Bio-Rad) and transferred into nitrocellulose or Immobilon®P membranes. Membranes were subsequently incubated with the indicated primary antibodies followed by the appropriate horseradish peroxidase-labelled antibodies (1/10000, Sigma-Aldrich or GE Healthcare, see Supplementary table). Immunodetection was performed using Super Signal West Pico or Femto Chemiluminescence Kit (Pierce). Quantification was done using Fiji software (Schindelin et al., 2012)).

### Molecular combing and detection by fluorescent antibodies

DNA was extracted and combed as described (Marheineke et al., 2009). Biotin was detected with AlexaFluor594 conjugated streptavidin followed by anti-avidin biotinylated antibodies. This was repeated twice, then followed by mouse anti-human ssDNA antibody, AlexaFluor488 rabbit anti-mouse, and AlexaFluor488 goat anti-rabbit for enhancement (Gaggioli et al., 2013). Images of the combed DNA molecules were acquired and measured as described (Marheineke et al., 2009). The fields of view were chosen at random. Several hundred of DNA fibers were analyzed for each experiment. Measurements on each molecule were made using Fiji software (Schindelin et al., 2012) and compiled using macros in Microsoft Excel. Replication eyes were defined as the incorporation tracks of biotin–dUTP. Replication eyes were considered to be the products of two replication forks, incorporation tracks at the extremities of DNA fibers were considered to be the products of one replication fork. Tracts of biotin-labelled DNA needed to be at least 1 kb to be considered significant and scored as eyes. When the label was discontinuous, the tract of unlabeled DNA needed to be at least 1 kb to be considered a real gap. The replication extent was determined as the sum of eye lengths divided by the total DNA length. Fork density was calculated as the total DNA divided by the total number of forks. The midpoints of replication eyes were defined as the origins of replication. Eye-to-eye distances (ETED), also known as inter-origin distances, were measured between the midpoints of adjacent replication eyes. Incorporation tracks at the extremities of DNA fibers were not regarded as replication eyes but were included in the determination of the replication extent or replicated fraction, calculated as the sum of all eye lengths (EL) divided by total DNA. Scatter plots of ETED and EL were obtained using GraphPad version 6.0 (La Jolla, CA, USA). Statistical analyses of repeated experiments have been included as means or medians including standard deviations or ranks as indicated in the legends. A p value**≤**0.05 was considered significant.

### Immunostaining and EdU labelling

For immunostaining, tadpoles were anesthetized in 0.005% benzocaine (Sigma), fixed in 1X PBS, 4% paraformaldehyde 1h at room temperature and dehydrated, then embedded in paraffin and sectioned (12 µm) with a Microm HM 340E microtome (Thermo Scientific). Immunostaining on retinal sections was performed using standard procedures. For proliferative cell labelling, tadpoles were injected intra-abdominally, 1-hour prior fixation, with 50-100 nl of 1 mM 5-ethynyl-2’-deoxyuridine (EdU, Invitrogen) at stage 41. EdU incorporation was detected on paraffin sections using the Click-iT EdU Imaging Kit according to manufacturer’s recommendations (Invitrogen).

Fluorescent images were taken with the AxioImagerM2 with Apotome (Zeiss) coupled to digital AxiocamMRc camera (Zeiss) and processed with the Axio Vision REL 7.8 (Zeiss) and Adobe Photoshop CS4 (Adobe) software. For quantifications of labelled cells by manual cell counting in the CMZ, a minimum of 16 retinas were analyzed. All experiments were performed at least in duplicate.

### Co-Immunoprecipitation

Immunoprecipitations from HEK293T cells expressing either HA-or FLAG-tagged Yap were performed using the Dynabeads Protein A Immunoprecipitation Kit (Invitrogen) by coupling 5 μg of anti-FLAG (Cell signaling) to the beads and following the manufacturer’s protocol. Immunoprecipitations from *Xenopus* egg extracts were performed as described below for mass spectrometry using rabbit anti-Yap (ab62752, Abcam) or rabbit anti-RIF antibodies. The 7.5% polyacrylamide gel was further analyzed by western blot using Mouse anti-Yap or rabbit anti-RIF antibodies Antibodies used for immunoprecipitation are listed in Supplementary table.

### Mass spectrometry

Rabbit anti-Yap antibody (ab62752, Abcam) or rabbit IgG (M7023, Sigma) were coupled 2 h at RT to protein A Sepharose beads (GE Healthcare). Coupled beads were covalently crosslinked using dimethyl pimelimidate according to standard procedures, washed with PBS and kept in PBS, 0.02% sodium azide at 4°C. For IP experiments, crosslinked beads with rabbit anti-Yap antibody or rabbit IgG were washed three times in EB buffer (50 mM Hepes, pH 7.5, 50 mM KCl, 5 mM MgCl2) and were incubated in *Xenopus* egg extracts for 30 min at 4°C. Beads were isolated by centrifugation and washed three times with EB buffer and once in EB buffer, 0.01 % Tween 20. The immunoprecipitated proteins were eluted by 2X Laemmli buffer and collected after centrifugation. Approximately 20 ng of immunoprecipitated Yap protein fraction was loaded on a 7.5% polyacrylamide gel and analyzed by mass spectrometry. Approximately 20 ng of immunoprecipitated Yap protein fraction was loaded on a 7.5% polyacrylamide gel and analyzed by mass spectrometry (Protéomique Paris Saclay-CICaPS platform). Protein samples were reconstituted in solvent A (water/ACN [98: 2 v/v] with 0.1% formic acid) and separated using a C18-PepMap column (Thermo Fisher Scientific) with a solvent gradient of 2–100% Buffer B (0.1% formic acid and 98% acetonitrile) in Buffer A at a flow rate of 0.3 µl/min. The peptides were electrosprayed using a nanoelectrospray ionization source at an ion spray voltage of 2300 eV and analyzed by a NanoLC-ESI-Triple TOF 5600 system (AB Sciex). Protein identification was based on a threshold protein score of > 1.0. For quantitation, at least two unique peptides with 95% confidence and a p-value < 0.05 were required.

Comprehensive protein list analysis and enriched biological pathways were based on Gene ontology classification system using Metascape (Sajgo et al., 2017). Data visualization was done using GOPlot R package (Livak and Schmittgen, 2001).

### Quantification and statistical analyses

For quantifications of labeled EdU^+^ cells by manual cell counting in the CMZ, 16 to 11 retinas per conditions with a minimum of 2 sections per retina were analyzed. Dissected eye areas and the number of cells per embryo were measured using Adobe Photoshop CS4 software. All experiments were performed at least in duplicate. Shown in figures are results from one representative experiment unless specified.

Statistical analyses (GraphPad Prism software, version 8.3.0) were performed using a Mann-Whitney test or Wilcoxon signed ranked test as mentioned in the figure legends.

### Data availability

The mass spectrometry proteomics data sets have been deposited to the ProteomeXchange Consortium via the PRIDE (Perez-Riverol et al., 2019) partner repository with the dataset identifiers PXD029345 and 10.6019/PXD029345.

## Acknowledgements

We are thankful to A. Kumagai and B. Dunphy for the gift of the C-terminal *Xenopus* Rif1 containing plasmid and *Xenopus* Rif1 antibody. We are grateful to A. Donval for her help with frog fertilizations and Virginie Chiodelli for technical support. This research was supported by grants to M.P. from ARC (Association pour la Recherche sur le Cancer), Association Retina France, Fondation Valentin Haüy, and UNADEV (Union Nationale des Aveugles et Déficients Visuels) in partnership with ITMO NNP (Institut Thématique Multi-Organisme Neurosciences, sciences cognitives, neurologie, psychiatrie) / AVIESAN (alliance nationale pour les sciences de la vie et de la santé). RMG was a Conacyt fellow (grant number 439641). This work has benefited from the facilities and expertise of the I2BC proteomic platform (Proteomic-Gif, SICaPS) supported by IBiSA, Ile de France Region, Plan Cancer, CNRS and Paris-Sud University.

---

**Figure 1 – figure supplement 1.**
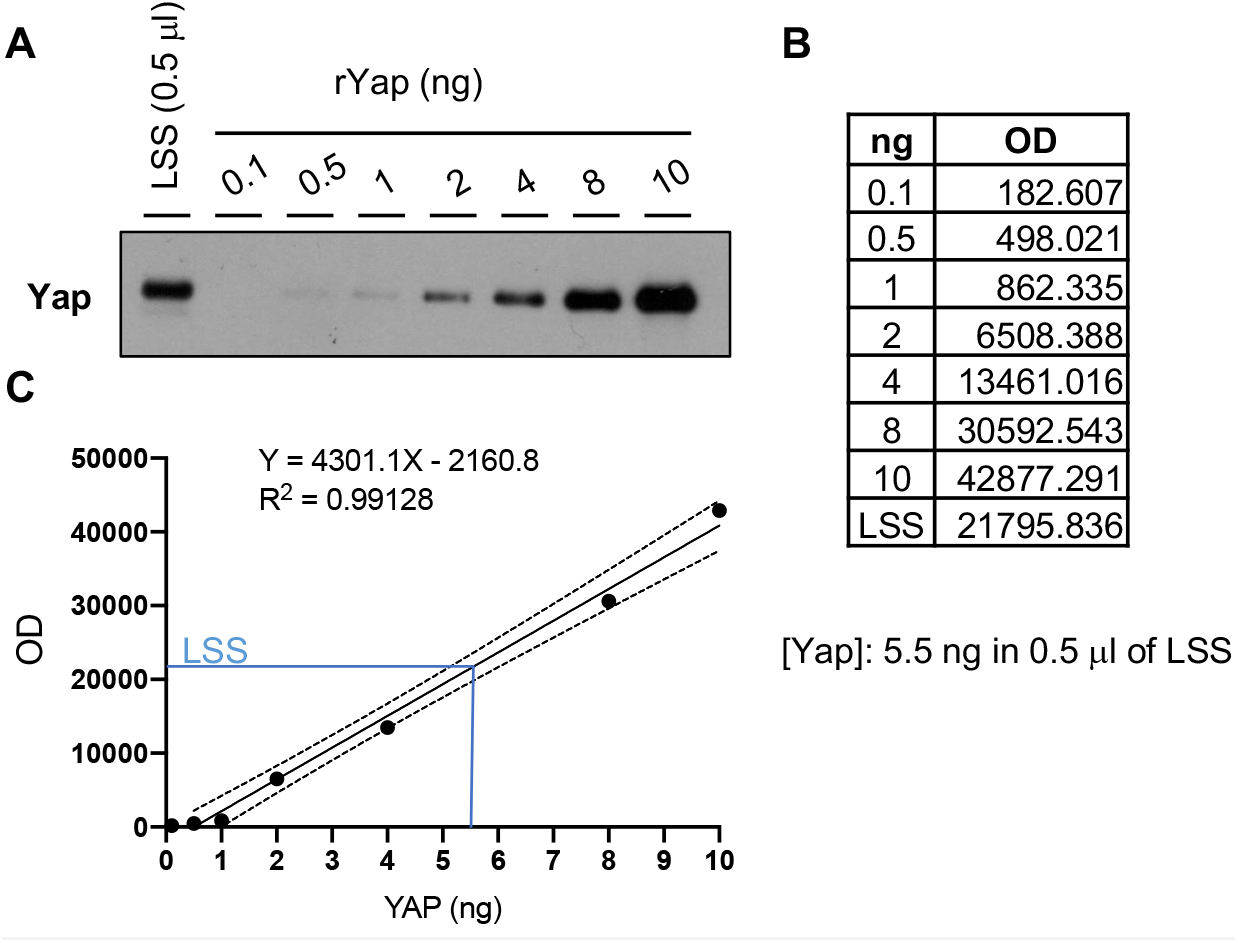
Yap protein concentration in *Xenopus* egg extracts. (**A**) Western blot showing different amounts of recombinant Yap (rYap) used to estimate endogenous Yap in egg extracts (low-speed supernatant, LSS). (**B, C**) The optical densities (OD) of the protein bands from (A) were used to plot a standard curve and to calculate Yap amount in the LSS.

**Figure 1 - figure supplement 2.**
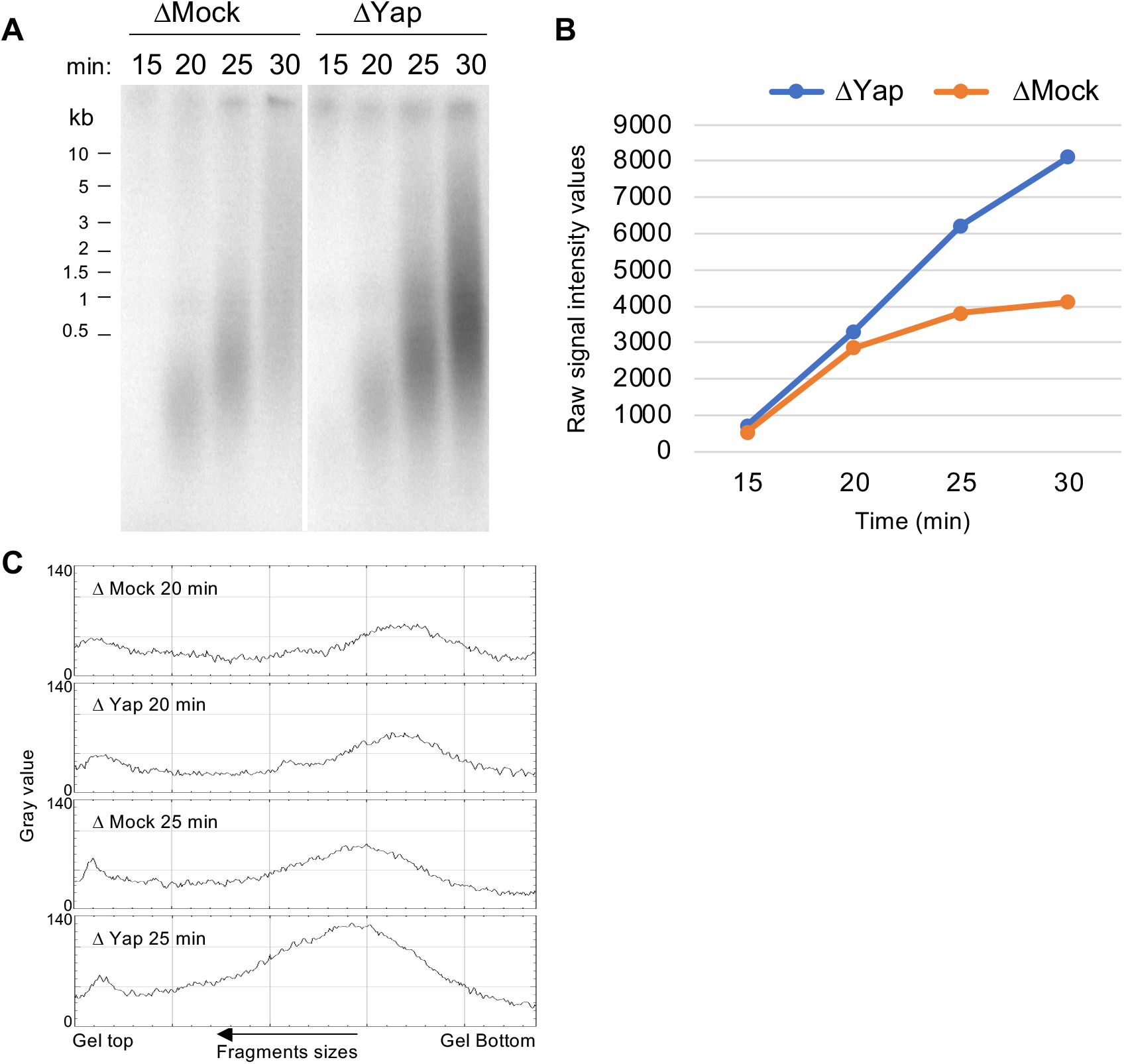
Yap depletion does not affect entry into S-phase. (**A**) Nascent DNA strands synthesized during early S-phase were analyzed at the indicated times by alkaline gel electrophoresis. (**B**) The amount of radioactivity incorporation was quantified for each lane and plotted as raw intensity values. Very similar incorporation is observed at the earliest time points (15-20 min) before getting higher in Yap depleted (ΔYap) compared to control depleted extracts (ΔMock) (25-30 min). (**C**) Grey scale profile (ImageJ) of lanes at 20 and 25 min, showing that size distribution of nascent strands is nearly identical in both conditions, and therefore entry in S-phase is unchanged after Yap depletion.

**Figure 1 - figure supplement 3.**
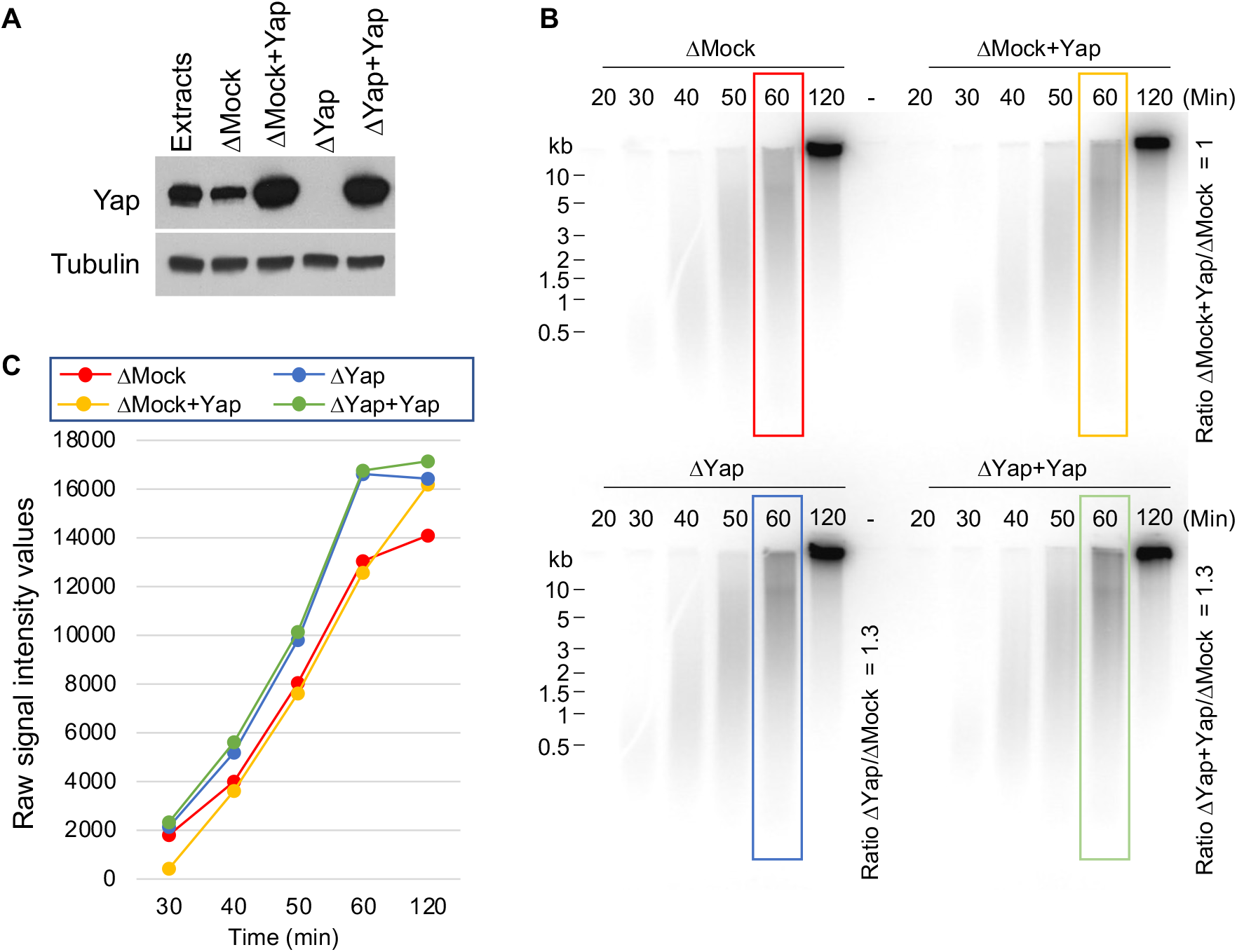
Adding back recombinant Yap protein does not restore the rate of DNA synthesis in Yap-depleted extracts. (**A**) Western blot showing the levels of Yap in either wild type egg extracts (Extracts), depleted egg extracts (ΔMock and ΔYap) or supplemented with *Xenopus* Yap recombinant protein produced in insect cells (ΔMock+Yap and ΔYap+Yap). (**B, C**) Nascent DNA strands synthesized were analyzed by alkaline gel electrophoresis after the indicated times (B). The level of radioactivity incorporation was quantified for each lane and plotted as raw intensity values (C). Ratios of the signal obtained in each condition over that of the ΔMock at 60 minutes is indicated in (B) as examples like in Figure 1D. -, unloaded lane.

**Figure 2 - table supplement 1.**
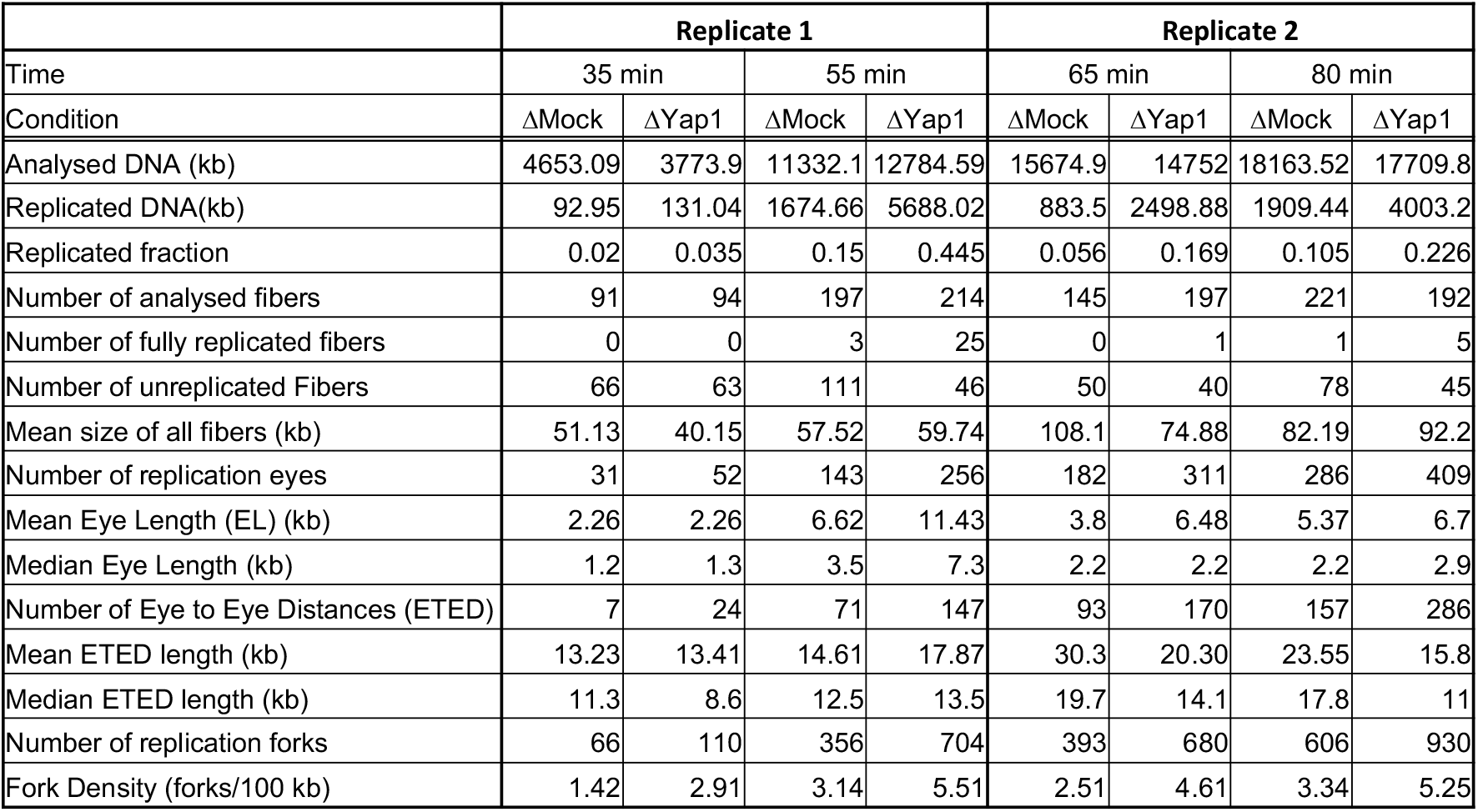
Depletion of Yap increases replication origin firing. Raw analysis of two independent DNA combing experiments presented in Figure 2. The analysis was performed as described in Materials and Methods.

**Figure 4 – figure supplement 1.**
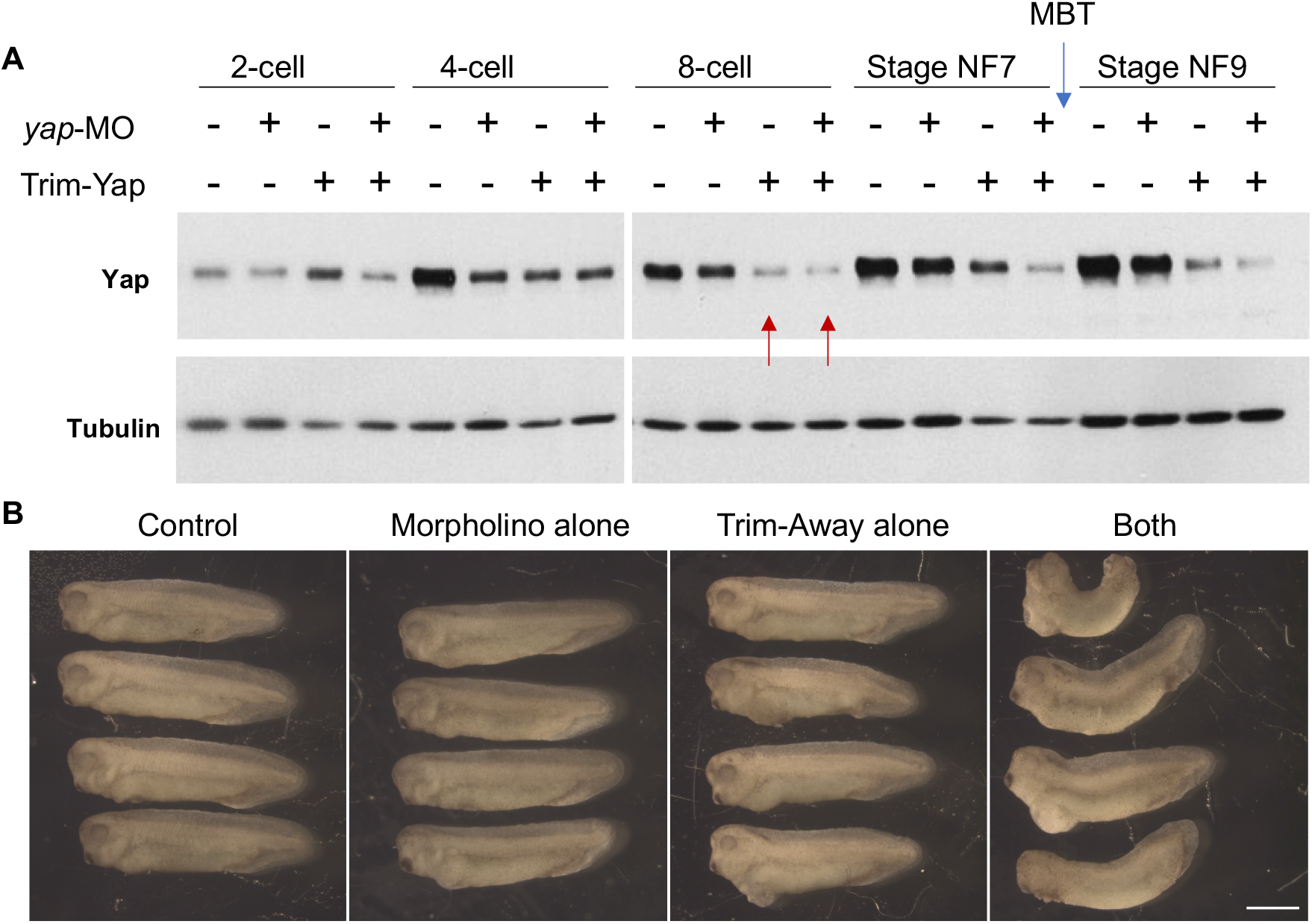
*Yap* knockdown using the Trim-Away strategy is effective at very early stages of development and leads to severe developmental defects in tadpoles. (A)Western blot performed on whole protein extracts from a pool of 8 embryos injected with either control morpholinos (-) or *yap* morpholinos (*yap*-MO, +), and/or pre-immune serum + TRIM21 (-) or anti-Yap antibodies + TRIM21 (Trim-Yap, +). Embryos were injected as described in Figure 4A then harvested at different times during development as indicated. The mid-blastula transition (MBT, blue arrow) is indicated as well as the time at which Yap depletion becomes observable (red arrows). (**B**) Injected embryos with (i) control MO + pre-immune serum + TRIM21 (Control), (ii) *yap*-MO + pre-immune serum + TRIM21 (Morpholino alone), (iii) control MO + anti-Yap + TRIM21 (Trim-Away alone) or (iv) *yap*-MO + anti-Yap + TRIM21 (both), were allowed to develop until the tadpole stage (NF32-33). Pictures of 4 specimens are shown for each condition. Scale bar = 1mm.

**Figure 5 – figure supplement 1.**
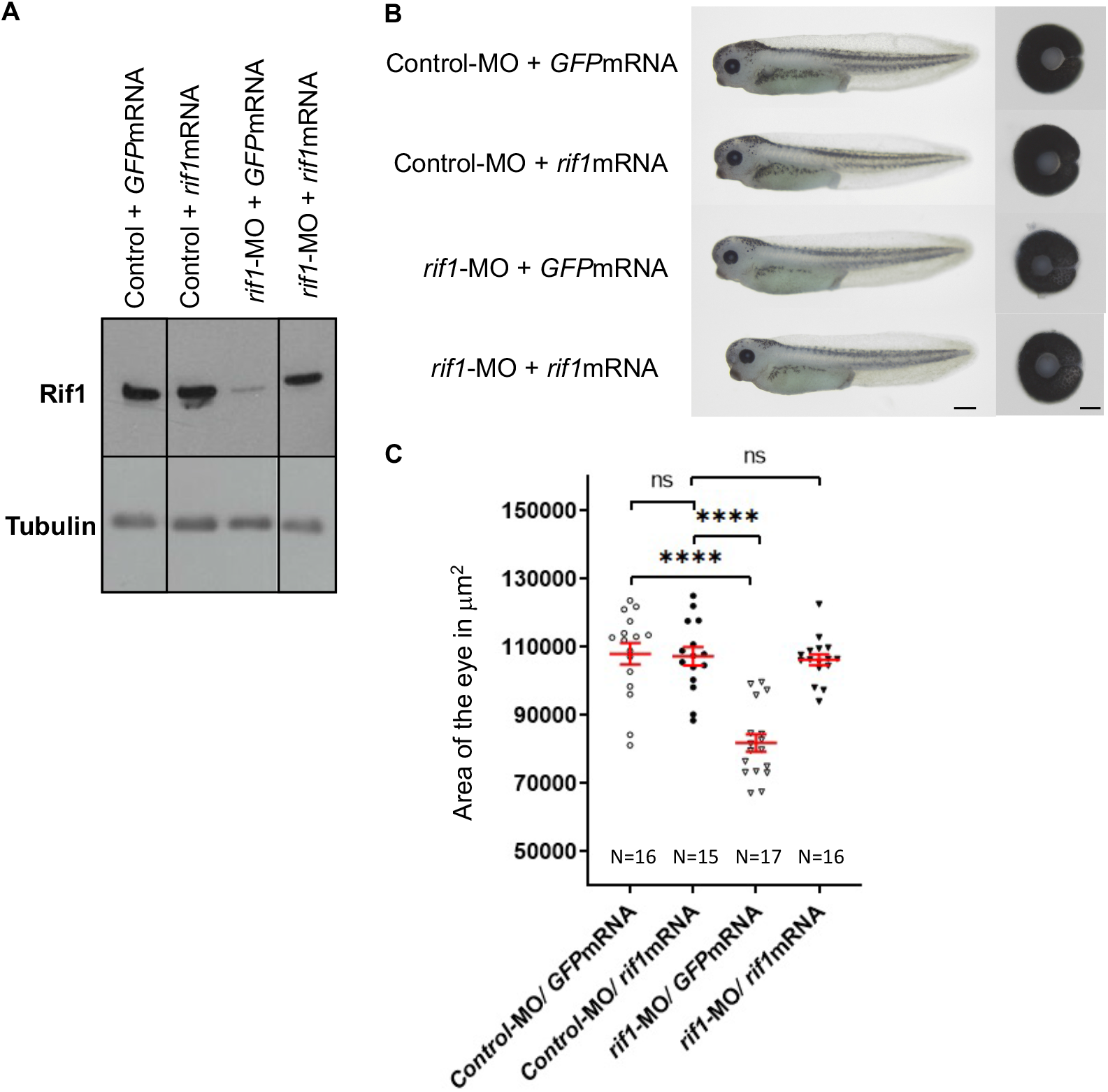
The *rif1*-MO-induced small eye phenotype is rescued by co-injection with *rif1* mRNA. (**A**) Western blot analysis showing the levels of Rif1 proteins in tadpoles at stage 41 following microinjection at 1 cell-stage of either Control-MO (Control) or *rif1*-MO together with GFP mRNA used as a control or *rif1* mRNA. Tubulin is used as a loading control. (**B**) Lateral views and dissected eyes of stage 41 tadpoles following one-cell stage microinjection of MO and mRNA as indicated. (**C**) Quantification of dissected eye areas. The *rif1*-MO-induced small eye phenotype is rescued by co-injection of *rif1* mRNA. Of note a suboptimal dose of *rif1* mRNA was used for the rescue experiment so that it does not alone generate any eye phenotype. The number of analyzed tadpoles is indicated for each condition. Data are shown as individual value plots with error bars (mean with SEM in red). Scale bar = 500 µm for tadpoles and 50 µm for dissected eyes.

## Notes

### Competing Interest Statement

The authors have declared no competing interest.

